# Finding rearrangements in nanopore DNA reads with last and dnarrange

**DOI:** 10.1101/2022.05.30.494079

**Authors:** Martin C. Frith, Satomi Mitsuhashi

## Abstract

Long-read DNA sequencing techniques such as nanopore are especially useful for characterizing complex sequence rearrangements, which occur in some genetic diseases and also during evolution. Analyzing the sequence data to understand such rearrangements is not trivial, due to sequencing error, rearrangement intricacy, and abundance of repeated similar sequences in genomes.

The last and `dnarrange` software packages can resolve complex relationships between DNA sequences, and characterize changes such as gene conversion, processed pseudogene insertion, and chromosome shattering. They can filter out numerous rearrangements shared by controls, e.g. healthy humans versus a patient, to focus on rearrangements unique to the patient. One useful ingredient is `last-train`, which learns the rates (probabilities) of deletions, insertions, and each kind of base match and mismatch. These probabilities are then used to find the most likely sequence relationships/alignments, which is especially useful for DNA with unusual rates, such as DNA from *Plasmodium falciparum* (malaria) with ∼ 80% `a`+`t`. This is also useful for less-studied species that lack reference genomes, so the DNA reads are compared to a different species’ genome. We also point out that a reference genome with ancestral alleles would be ideal.

## 1 Introduction

The last software was first made public in 2008: it was intended as a general tool for finding and aligning related regions in gigabase-scale sequence data. It is not really designed for recent long-read data such as nanopore, nevertheless it can be used, and has some unique advantages. The main advantages are:

1. It can learn the rates (probabilities) of deletions, insertions, and each kind of base match and mismatch (Fig. 1), which are due to a combination of sequencing errors and real sequence differences. It then uses these probabilities to determine the most probable alignments (*1*).

**Figure 1:**
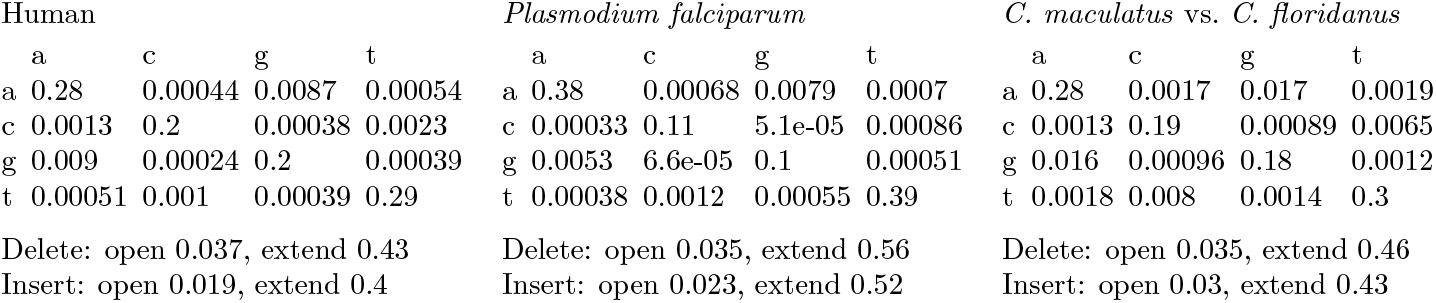
Rates (probabilities) of base matches, mismatches, deletions, and insertions between nanopore DNA reads and reference genomes. In each 4×4 matrix, a row corresponds to a genomic base, and a column to a read base.
2. It can disentangle complex rearrangements and duplications between sequences, even when confounded by repeats, by finding the most probable split of a sequence into rearranged parts based on the alignment probability of each part (*2*).

Another useful feature is that last can compare DNA to protein sequences, allowing frameshifts, which are frequent in long DNA reads. This has been used for taxonomic and functional binning of metagenomic long reads (*3*).

Here we shall focus on finding rearrangements from last alignments of nanopore DNA reads to a reference genome. The next problem is that we typically find thousands of rearrangements, many of which seem to be artifactually-rearranged reads, and many others real-but-benign (*4, 5*). The artifacts seem to be sporadic, so they can be depleted by sequencing to several-fold coverage of the genome and then discarding reads with unique rearrangements not shared by any other read (*4, 5*). When seeking rearrangements causing genetic disease, we also use nanopore reads from control individuals without the same disease, and discard benign rearrangements shared with controls. These tasks downstream from last are done by `dnarrange`.

### 1.1 Examples of match, mismatch, insertion, and deletion rates

Three examples of these rates, for nanopore DNA reads versus reference genomes, are shown in Fig. 1. The human rates are for one set of human nanopore reads, `HG02723_1` (*6*), versus a reference human genome (hg38). We can see, for example, that `a`↔`g` substitutions are much more frequent than other substitutions, which is often the case for nanopore sequences (*4, 5*). Also, the probability (rate) of opening a deletion is much higher than an insertion. The *P. falciparum* rates are for one set of nanopore reads (*7*) versus a reference genome: these are interesting because *P. falciparum* DNA is ∼ 80% `a+t`, so the match and mismatch rates are quite different from human. The final example is a set of nanopore reads from the ant *Camponotus maculatus* (*7*) versus the closest-available reference genome: *Camponotus floridanus*. This example has higher substitution rates, as expected between different species.

### 1.2 Understanding rearrangements

Various mutational processes can cause almost arbitrarily complex rearrangements, duplications, and deletions of DNA sequence (Fig. 2). For example, chromosomes can shatter into many fragments that rejoin in a different order and orientation, or DNA polymerase can template-switch during DNA replication causing rearrangements, duplications, and deletions. No matter how complex these changes, there is one key point: almost every part of a descendant sequence comes from a unique part of ancestral sequence(s). This is shown in Fig. 2b: we can visually scan the derived sequence from top to bottom and see where each part comes from (joined-up diagonal lines).

**Figure 2:**
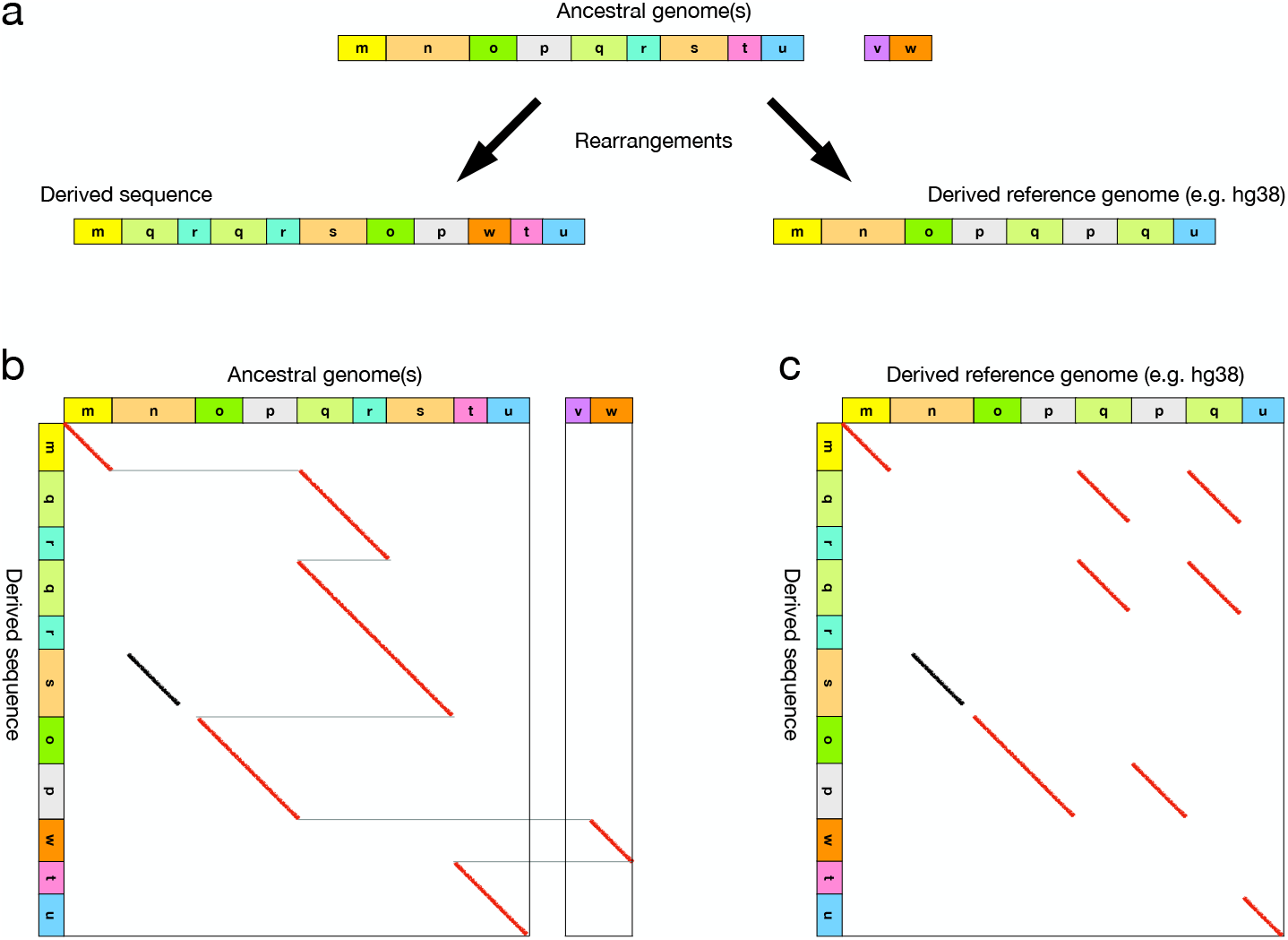
A made-up example of complex rearrangements during DNA sequence evolution. Each colored block represents a segment of DNA, e.g. a few hundred bp. These segments become rearranged, duplicated, or deleted. The blocks labeled n and s are similar sequences from an earlier duplication, i.e. paralogs. Reproduced from (*5*).

We might wonder about “insertions”: where does the inserted sequence come from? It must come from somewhere: typically it has been duplicated or moved from elsewhere in the genome. Rarely, an insertion may come from a different genome, such as a virus. There is also non-templated insertion aka “spontaneous generation” of new sequence with no ancestor: this is the only exception to the rule.

Unfortunately we do not have ideal ancestral sequences: in practice we compare DNA reads to a “reference” genome which has separately undergone complex changes. The relationship between two descendant sequences is fundamentally less simple (Fig. 2c), e.g. duplication in both lineages creates a many-to-many relationship, and reference-specific deletion means that parts of DNA reads correctly align nowhere in the reference, but may align incorrectly to paralogs. Even if we could perfectly detect the correct relationships (diagonal red lines in Fig. 2c), it is hard to understand what changes have occurred.

In practice, we assume the reference genome is ancestral, even though it is not. We are interested in rearrangements that cause genetic disease: at least in these cases the reference is likely to represent the ancestral state. If virus insertions are suspected, viral chromosomes can be added as extra chromosomes to the reference. We will find spurious rearrangements because the reference is not perfectly ancestral: the hope is that these will also be found in control data, thus discarded. Another way to detect reference-genome regions with non-ancestral status is by comparing to an outgroup, e.g. an ape genome (*4*). It would be useful to have a reference genome that contains ancestral alleles (*4, 5*).

### 1.3 Simple sequences

DNA is rife with “simple sequences” such as catcatcatcat or aaaattaaaacaaa. They cause many similarities between sequences that are not correct relationships, because they are not descended from a common ancestor. There are several methods for detecting and “masking” simple sequences, which do not all work equally well (*8*). last uses `tantan` (*8*) to detect simple sequences, and converts them to lowercase letters. However, masking can cause problems by hiding correct relationships between sequences, which might cause incorrect relationships to be found instead. So, by default last treats lowercase the same as uppercase when finding alignments. Alignments without a significant amount of uppercase- to-uppercase alignment are suspicious, and can optionally be discarded by `last-postmask`.

## 2 Methods

### 2.1 Installation

It may be easiest to install the software from Bioconda (*9*), or from Debian Med (*10*). After setting up Bioconda on your computer, this command installs `dnarrange` and all its dependencies, including last (see Note 1):

~~~
conda install dnarrange
~~~

The following discussion applies to last version ≥ 1387 and `dnarrange` 1.5.2, and not to older versions.

### 2.2 *Getting the* Camponotus *rates*

Let us first see how to get the ant results in Fig. 1. We need the reference genome sequence in fasta format, which we got by searching “camponotus” at NCBI Genome (https://www.ncbi.nlm.nih.gov/genome/). We renamed this file `Cflo_v7.5.fa`. The first step is to prepare “index” data-structures for the genome, which enable fast sequence comparison:

~~~
lastdb -P16 -uNEAR antDB Cflo_v7.5.fa
~~~

This creates several files whose names start with `antDB`. The `-P16` option makes it faster by running 16 parallel threads, with no effect on the result. Adjust as appropriate for your computer (see Note 2).

The `-uNEAR` option changes the seeding scheme. LAST uses a seed-and-extend heuristic: it first find “seeds”, simple gapless alignments, then tries to extend full alignments from the seeds. Although the seeds are gapless, they can allow some mismatches: the more mismatches they allow, the longer they need to be to retain specificity. `-uNEAR` specifies short seeds with few mismatches, which is appropriate for searching indel-rich nanopore reads against a closely-related genome. If you omit `-uNEAR`, it may not make much difference in practice.

Next, we need the DNA reads in fasta or fastq format. We do not use the extra information in fastq, so fasta is preferable because the files are much smaller. We can find the rates of substitution etc. between reads and genome like this, using the original fastq file in this case:

~~~
last-train -P16 -Q0 antDB CmacRNAseq_180528.fastq.gz > ants.train
~~~

The `-Q0` option makes it ignore the fastq quality data (see Note 3). If you have fasta instead of fastq, `-Q0` has no effect and can be omitted.

It’s worth noting that `last-train` doesn’t actually use all the DNA reads: it uses a random sample of size one million bases (probably overkill). This means that giving huge read files to `last-train` doesn’t make it much slower, except for the time needed to read the file and get a random sample.

`last-train` works iteratively: it compares the reads to the genome to find the rates, then uses these rates to do the comparison more accurately and get better rates, and so on. It prints data for each iteration (for troubleshooting), with the final rates at the end. It’s not necessary to look at them: you can just pass the output file to the next alignment step.

### 2.3 *Getting the* Plasmodium falciparum *rates*

The *P. falciparum* rates were found like this:

~~~
lastdb -P16 -uNEAR -R02 plaDB plaFal3D7.fa
last-train -P16 -Q0 plaDB P.falciparum_targeted_seq.fastq > pf.train
~~~

The only change is the `-R02` option, which is recommended for DNA with ∼ 80% `a+t`. This option changes the `tantan` parameters for defining “simple sequence”, which is necessary for very `(a+t)`-rich DNA. If you omit -R02, it might not matter too much in practice. `last-train` avoids lowercased simple sequence, but in this case it hardly affects the result.

### 2.4 Getting the human rates

The human rates were found like this:

~~~
lastdb -P16 -uRY4 hdb hg38.analysisSet.fa
last-train -P16 -Q0 hdb HG02723_1.fastq.gz > hum.train
~~~

The change here is to use `-uRY4` instead of `-uNEAR`. This makes last faster and use less memory, at a cost in sensitivity. The main reason for doing this is that the fastq file is quite big, 81 gigabases. This is not necessary for `last-train`, which just uses a sample of the fastq: the point is to speed up the subsequent alignment step. `RY4` makes it use ¼ of the seeds, in a similar way to this: just use seeds starting with a (see Note 4).

This `last-train` run took 23 minutes, most of which was spent decompressing the fastq file to get a random sample from it. It is not necessary to re-run `last-train` for each dataset, unless there is reason to think the rates may have changed, e.g. due to different versions of sequencing hardware or base-calling software.

### 2.5 Aligning human DNA reads to a human genome

The next step is to actually align all the reads to the genome:

~~~
lastal -P16 --split -p hum.train hdb HG02723_1.fastq.gz | gzip > out.maf.gz
~~~

This `lastal` command works in two stages. First, it finds and aligns similar regions of the reads and the genome, often aligning the same part of a read to several genome regions. Second, `--split` makes it cut these alignments down to a unique best alignment for each part of each read. This is appropriate if the genome is ancestral to the reads (Fig. 2). Sometimes, part of a read matches two or more genome regions with almost equal probability: `--split` choses the most probable alignment (arbitrarily if exactly equal), and outputs “mismap probabilities”, the probability that each part of a read is aligned to the wrong place. Finally, `gzip` compresses the output (optional, see Notes 5 and 6). This alignment took ∼ 6 hours, with 16 threads (`-P16`).

### 2.6 An alternative way using WindowMasker

Our published human studies have mostly not used RY4 (because it’s somewhat new), and instead saved time by “masking” repeats. The main cause of alignment slowness is the abundance of repeats, such as LINEs, SINEs, and simple sequences, so each repeat in a read gets preliminarily aligned to multiple genome locations.

We can mitigate this problem by masking repeats. We wish to mask as little as possible in order to maximize sensitivity, but enough to make the run time tolerable. For this aim, WindowMasker (*11*) seems to work well: it is part of the BLAST package, which can be got from NCBI, Bioconda, or Debian Med. The following commands find repeats in the genome and convert them to lowercase:

~~~
windowmasker -mk_counts -in hg38.analysisSet.fa > hum.wmstat
windowmasker -ustat hum.wmstat -outfmt fasta -in hg38.analysisSet.fa > hum-wm.fa
~~~

The next step is to run lastdb like this:

~~~
lastdb -P16 -uNEAR -R11 -c hwmdb hum-wm.fa
~~~

The `-R11` option retains lowercase from the input and additionally lowercases simple repeats found by `tantan`. The `-c` tells it to “mask” lowercase: this will exclude lowercase from seeds but not from final alignments, and then discard alignments that lack a significant amount of uppercase-to-uppercase alignment (the same as `last-postmask`). The subsequent `last-train` and alignment steps are the same as above.

### 2.7 Finding rearrangements with dnarrange

`dnarrange` gets rearranged DNA reads from the read-to-genome alignments, and performs two optional filtering steps. It discards reads with unique rearrangements not shared by any other read, and discards reads with rearrangements shared by control reads. So we need control DNA reads aligned to the same genome. A few control sets aligned to hg38 are available at https://github.com/mcfrith/dnarrange. We can run `dnarrange` like this:

~~~
dnarrange -v out.maf.gz : controls/hg38-* > groups.maf
~~~

The -v (verbose) option just makes it show progress messages as it’s working (useful for troubleshooting and reassurance that it’s doing something). The “:” separates case from control files (see Note 7). The output has the alignments of the rearranged reads, in groups where reads that cover the same rearrangement are in the same group. In our studies of human patients, the number of groups per patient declines from thousands without controls to a few dozen after control filtering (*5*). Filtering is more effective when some controls are from the same family or ethnic group as the case individual.

It’s possible to not use control DNA reads, but then an overwhelming number of rearrangements may be found. They may include dubious rearrangements in regions where the reference genome does not represent the ancestral state, or where the reference is incomplete. For example, if a segment of the reference has been deleted, the corresponding part of a DNA read may get wrongly aligned to a paralog elsewhere in the genome, showing an incorrect rearrangement.

An alternative is to use `last-postmask` (if repeats were not already masked with `lastdb` option `-c`):

~~~
last-postmask out.maf.gz | dnarrange -v - : controls/hg38-* > groups.maf
~~~

The “-” has a standard meaning of reading the data that is piped in. As mentioned above, `last-postmask` discards alignments that are mostly of lowercased simple sequence. There is doubt that such alignments reflect correct relationships. Nevertheless, such alignments can be informative because they do indicate changes such as expansion or insertion of simple sequence (see Note 8).

### 2.8 Making dotplot figures of the rearrangements

We can make a dotplot figure of each group in `groups.maf` like this:

~~~
last-multiplot -a refGene.txt -a rmsk.txt groups.maf fig-dir
~~~

This puts the figures in a new directory `fig-dir`. We have used the -a option to show genes and repeats, using files downloaded from the UCSC Genome Browser (http://genome.ucsc.edu). It’s also possible to show BED, GFF/GTF, or RepeatMasker .out data, or unsequenced gaps in AGP or gap.txt format (see Note 9). Naturally, these dotplots are more useful when there are not too many of them, and making many of them is slow.

Some examples of human rearrangements, in `HG02723_1` versus hg38, are shown in the remaining Figures. Fig. 3 shows a classic genetic phenomenon: gene conversion. This is a process where one DNA sequence gets replaced, probably during DNA repair, by a copy of a similar sequence. In this case, part of an L1 LINE element in chromosome 20 has been replaced by the homologous part of another L1 element. Here, the converted read parts are aligned to one L1 in chromosome 6, but their mismap probabilities are quite high (not shown), because they have almost equally probable alignments to other L1s. So the specific donor element may not be confidently knowable.

**Figure 3:**
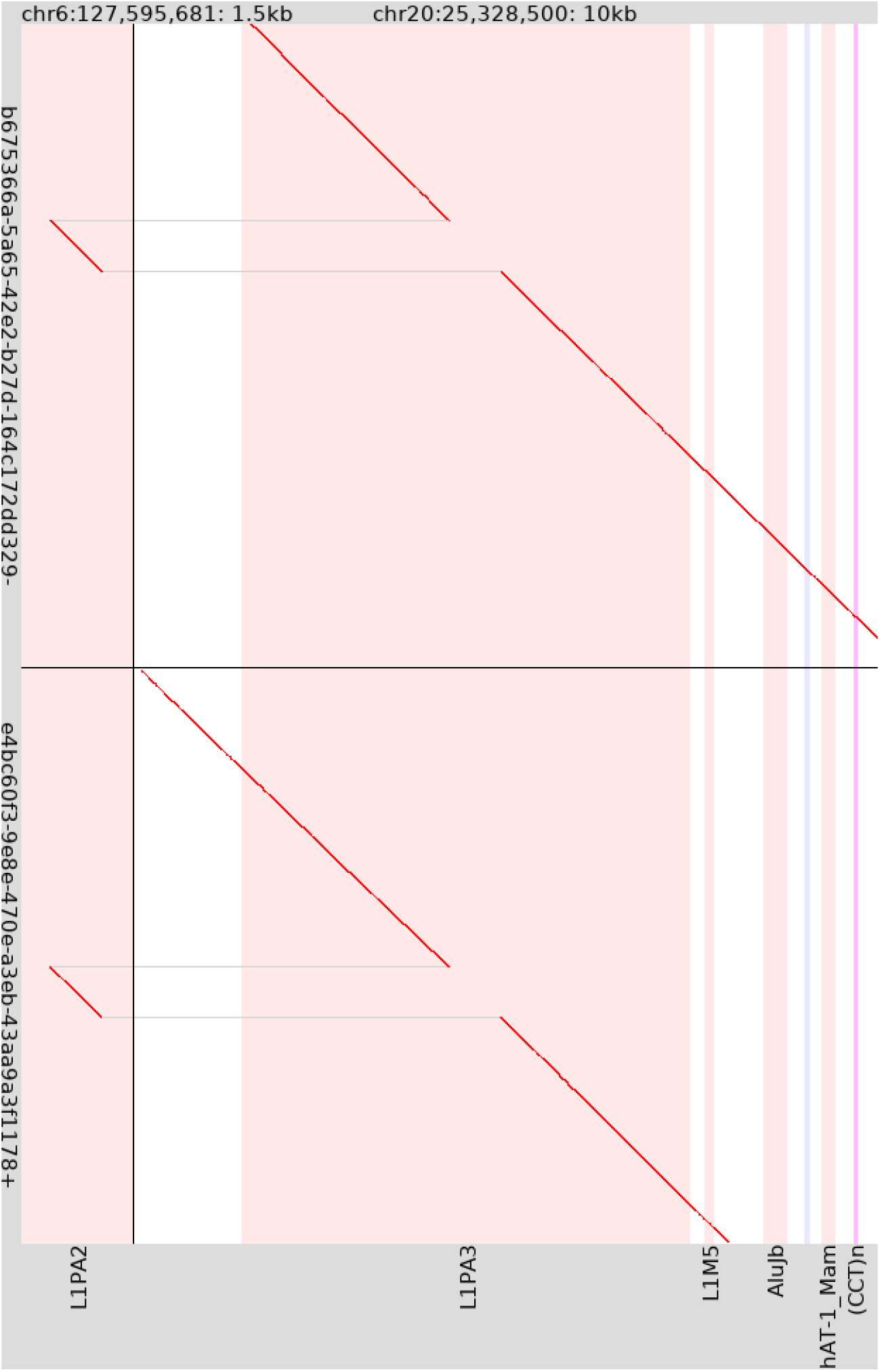
Alignment of 2 human DNA reads to human genome hg38, showing gene conversion. The read identifiers are shown on the left: the final − or + indicates that the read has been reverse-complemented or not. The figure shows only part of each read. To the left of the vertical black line is 1.5 kb of chromosome 6, to the right is 10 kb of chromosome 20. The vertical stripes show repeat elements in hg38. (The stripes have 3 different colors for forward-oriented elements, reverse-oriented elements, and simple repeats.)

Gene conversion is a poster child for probability-based sequence alignment of the sort done by `--split` (*4*). A naive alignment method would just align these DNA reads continuously to this L1 in chromosome 20, because the converted sequence is similar to the replaced sequence.

Fig. 4 shows a short-range rearrangement, localized within 5 kb, with a rather small rearranged fragment. This resembles mutations that have been attributed to template switching during DNA replication, which can create numerous complex mutation patterns, and have caused erroneous variant annotations (*12*).

**Figure 4:**
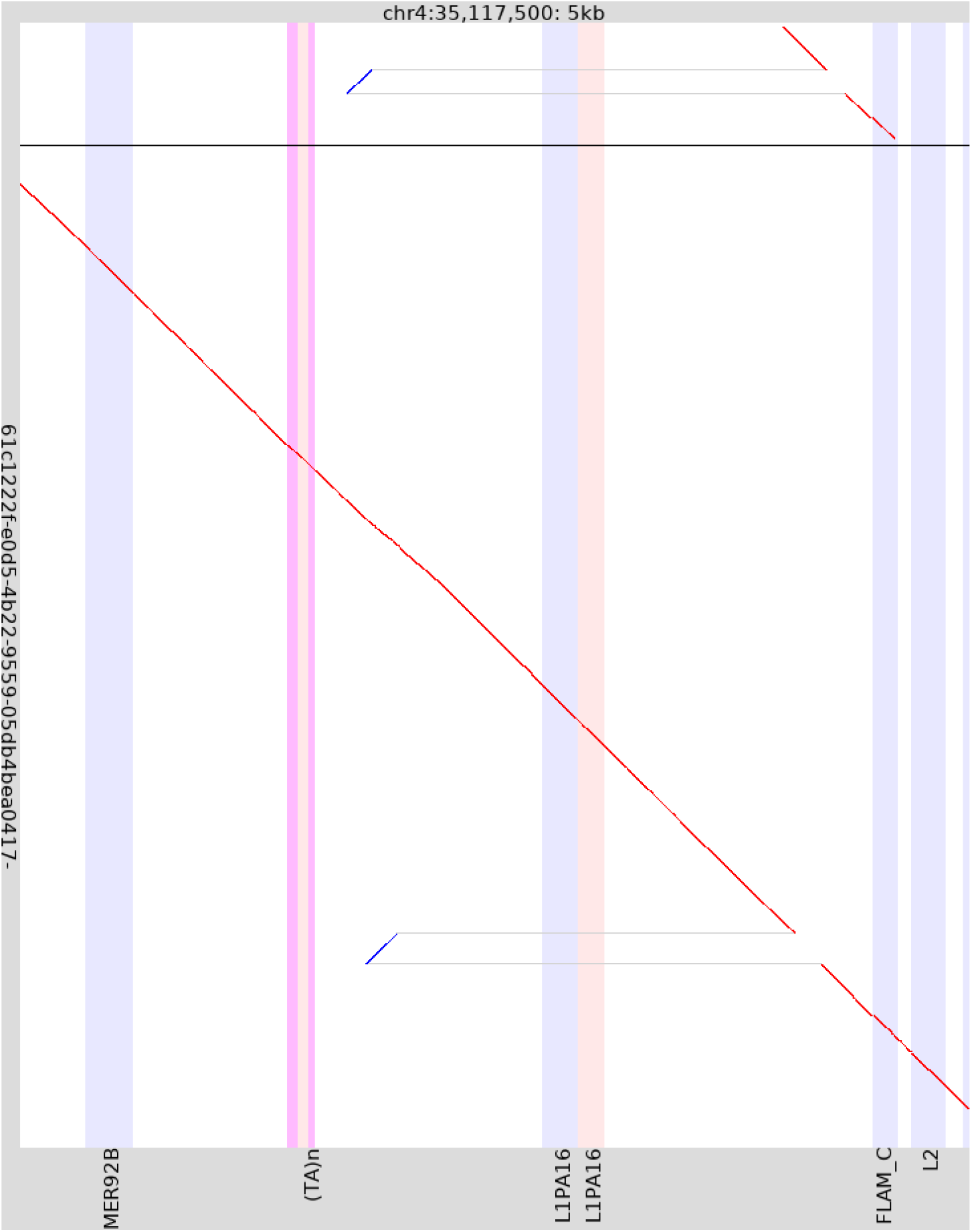
Alignment of 2 human DNA reads to human genome hg38, showing a short-range rearrangement. The top read is short, and its identifier is omitted.

Fig. 5 shows inertion of a processed pseudogene from chromosome 11 into chromsome 13. This is a process where an mRNA molecule, after its introns have been removed, is reverse-transcribed and inserted into the genome (perhaps by a reverse transcriptase enzyme from a retrotransposon).

**Figure 5:**
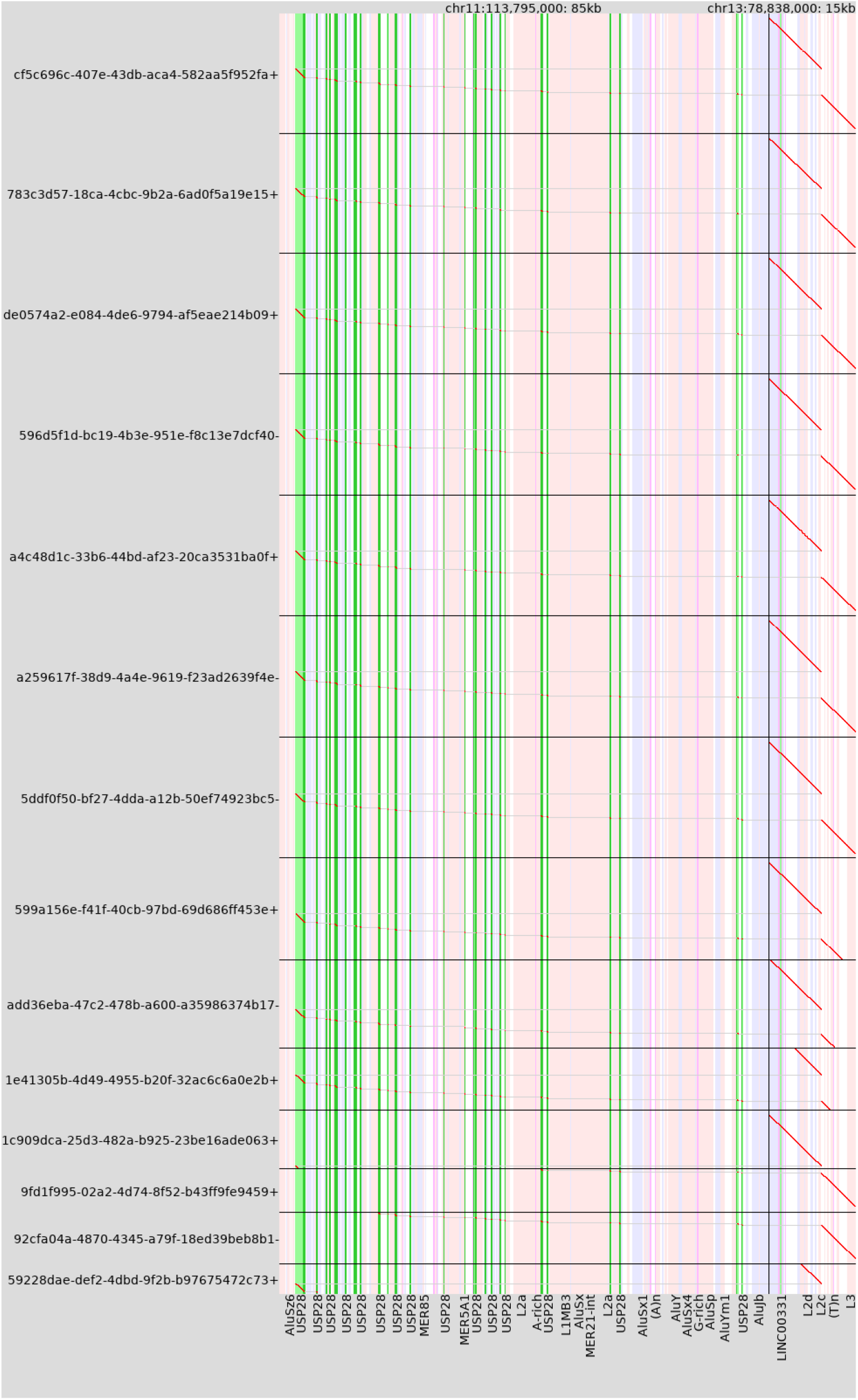
Alignment of 14 human DNA reads to human genome hg38, showing insertion of a processed pseudogene from chromosome 11 into chromosome 13. The vertical green stripes show exons of *USP28*, which encodes ubiquitin specific peptidase 28.

Finally, Fig. 6 shows a DNA read aligning to chromosome 1, with a gap in the alignment, which might be a deletion in the read or an insertion in the reference. The gap coincides exactly with an L1PA2 element, which is a young LINE retrotransposon. There is no known mechanism for precise excision of a LINE element, whereas LINE insertion is normal, so this is an insertion that occurred in the lineage leading to the reference sequence. This is an example where the reference does not have the ancestral state. Outgroups (other mammal genomes) lack this L1PA2 (Fig. 6).

**Figure 6:**
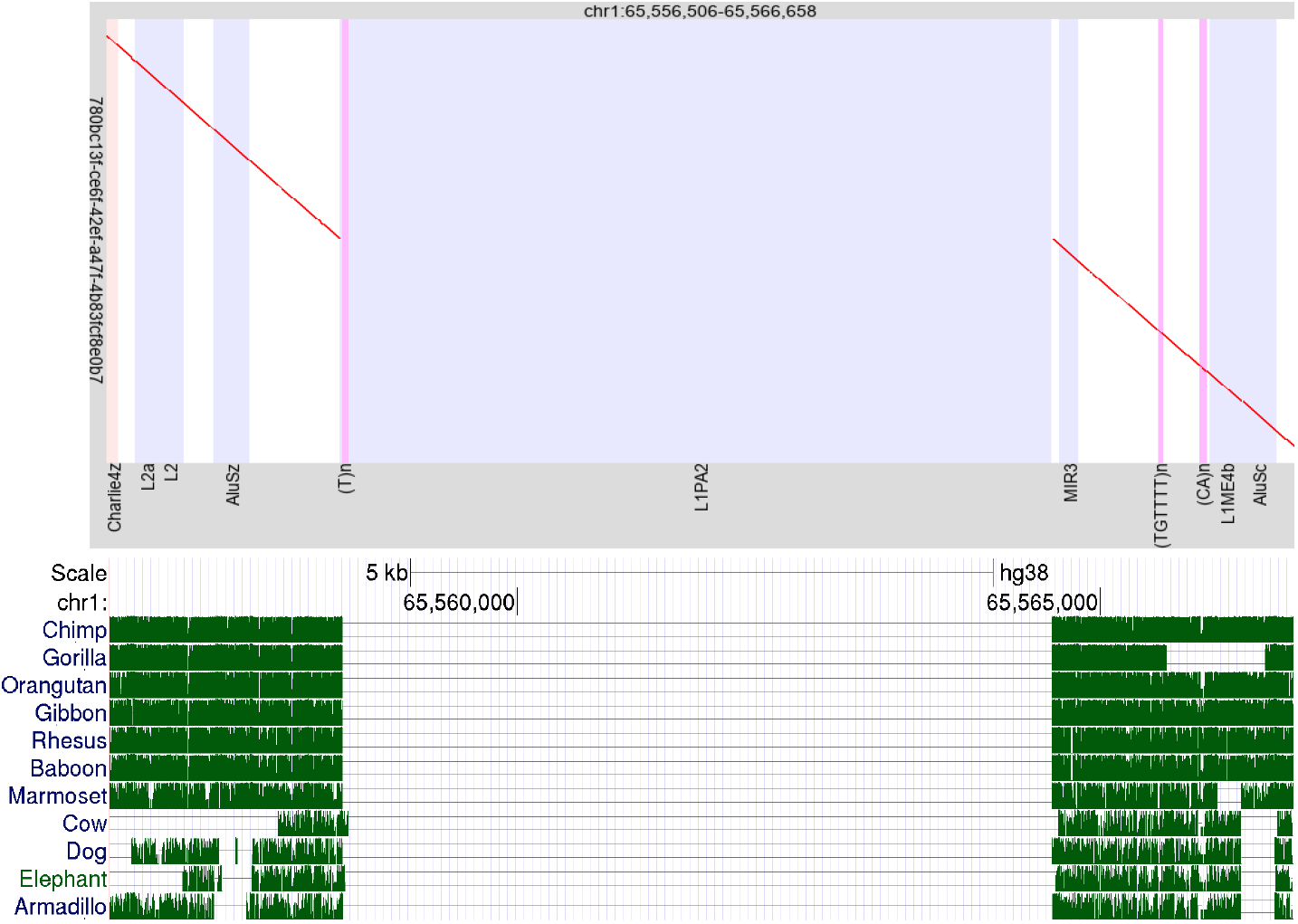
Above: alignment of a human DNA read to human genome hg38, showing reference-specific insertion of an L1 LINE retrotransposon in chromosome 1. Below: alignments of other mammal genomes to this part of chromosome 1 (screen shot from http://genome.ucsc.edu).

### 2.9 last-dotplot

`last-multiplot` (which comes with `dnarrange`) uses `last-dotplot` (part of last) to draw each figure. It may be useful to run `last-dotplot` directly. For example, Fig. 4 was made like this:

~~~
last-dotplot -a refGene.txt -a rmsk.txt -j2 --labels1=2
                                    -1 chr4:35117500-35122500 groups.maf fig4.png
~~~

Option -j2 draws the gray lines joining the aligned segments, `--labels1=2` adds the start coordinate and length to the top of the figure, and `-1 chr4:35117500-35122500` shows that range of `chr4`.

### 2.10 Rearrangement types and thresholds

A rearrangement is indicated when two parts of a DNA read align to disjoint places in the genome. `dnarrange` classifies four kinds of disjointness:

1. Different chromosomes (Fig. 3)
2. Opposite strands of one reference sequence (Fig. 4)
3. Non-colinear alignment of two read parts to the same strand of the same reference sequence. In other words: the alignment of the upstream read part ends at coordinate *X* in the reference sequence, the alignment of the downstream read part starts at coordinate *Y*, and *Y* is upstream of *X*.
4. Two consecutive read parts (i.e. with no aligned part between them in the read) that align colinearly with a big gap in the reference (Fig. 6).

By default, `dnarrange` ignores “big gaps” *<* 10 kb, and non-colinearities where *Y* is less than 1 kb upstream of *X*. This is because small deletions (colinear gaps) are frequent and arguably not “rearrangements”, and small tandem duplications (non-colinear) are overwhelmingly numerous. You can set the g (gap) and r (reverse jump) thresholds in bp:

~~~
dnarrange -v -g1000 -r100 out.maf.gz : controls/hg38-* > groups.maf
~~~

We have said that `dnarrange` discards DNA reads that share rearrangements with control reads, but the truth is a bit more complex. `dnarrange` prioritizes rearrangement types in this order: inter-chromosome *>* inter-strand *>* non-colinear *>* gap. It discards DNA reads that share rearrangements of the highest-priority type present in the read with control reads.

### 2.11 Other features of dnarrange

We have covered the most important features of `dnarrange`, but it can do some further things. `dnarrange-merge` merges each group of DNA reads into a consensus sequence, using `lamassemble` (*13*). By re-aligning the consenus sequence to the genome, we can perhaps see the rearrangement more clearly and accurately. More simply, `dnarrange-merge` can just get the rearranged reads without merging them:

~~~
dnarrange-merge HG02723_1.fastq.gz groups.maf > some-reads.fastq
~~~

This may be useful, because we can re-align just these reads to the genome more slowly and sensitively, for example like this:

~~~
lastdb -P16 -uNEAR nearDB hg38.analysisSet.fa
lastal -P16 -m50 --split -p hum.train nearDB some-reads.fastq > out2.maf
~~~

Here we have used `NEAR` instead of `RY4` for greater sensitivity. We have re-used the `last-train` result: no need to re-run it. Finally, we added option `-m50`, which makes it even more slow and sensitive. Higher m values make it increasingly slow and sensitive: the default is 10 (but the `-uRY` options set the default to 2). A further issue is that some large and complex rearrangements, e.g. in genetic diseases, are larger even than “long” reads. So each group of reads covers only part of the rearrangement. In order to understand the whole rearrangement, we need to correctly order and orient the groups (i.e. the rearrangement parts). `dnarrange` includes a method that seeks a most-parsimonious order and orientation, which succeeded in fully characterizing some large and complex rearrangements (*5, 14*). Interestingly, these rearrangements have holistic features, e.g. loss of sequence, that are knowable only from the whole rearrangement and not from the parts.

`dnarrange` can be very slow and memory-consuming. This seems to be because many DNA reads align dubiously to a few genomic hotspots, and `dnarrange` spends much effort comparing these reads to each other. The solution is to use control reads, which rapidly discard these hotspot reads. This works better if the case and control reads were analysed in the same way, e.g. with/without masking. It also works better if the exact same reference was used. (There are unfortunately different versions of e.g. hg38). Another way to mitigate this problem is to use `last-postmask`.

It is useful if the DNA reads are long enough to cover a whole rearrangement, but on the other hand long reads have a disadvantage. The problem is that if a read overlaps two rearrangements, and one of them is shared by controls, the whole read may get discarded. This is a limitation of `dnarrange`, because it is hard to know whether a read covers two small rearrangements or two parts of one large rearrangement. Therefore, a combination of longer and not-so-long reads may be best.

Our previous publications show further interesting rearrangements found by these methods, such as tandem heptuplication, 3’-transduction from a LINE, or chromsosome shattering (*4, 5*).

## 3 Notes

1. At the time of writing, `conda` seems to have a bug where it may install old versions of software: it may be better to use `mamba` (https://github.com/mamba-org/mamba).
2. As a special case, -P0 uses as many threads as your computer claims it can run simultaneously, which is a good way to annoy the other users of a shared server.
3. last can alternatively use the quality data for more accurate training and alignment. This assumes, however, that the qualities indicate probability of substitution error, not insertion or deletion error, and we are not confident this holds for nanopore data.
4. Actually, `RY4` uses seeds starting with combinations of purines and pyrimidines (*15*). There are faster alternatives using 1⁄8, 1⁄16, and 1⁄32 of the seeds: `RY8`, `RY16`, and `RY32`.
5. We can use `gzip` options to get faster but worse compression. For example, `gzip -5` shaves ∼ 20% off the alignment run time, and adds 6% to the output size.
6. The data transfer from `lastal` to `gzip` is inefficient, because `lastal` produces output in bursts, and `gzip` takes time to absorb each burst, making lastal wait. This can be fixed with `mbuffer`: `lastal -P16 --split -p hum.train hdb HG02723_1.fastq.gz | mbuffer | gzip > out.maf.gz mbuffer` absorbs the bursts quickly, and feeds them to `gzip. mbuffer` is available in Bioconda at the time of writing, although it is not biology specific.
7. It’s possible to give `dnarrange` more than one case file: it will only output groups that have reads from all case files.
8. To analyze tandem repeat changes, we have specialized software `tandem-genotypes` (*16*), described elsewhere in this volume.
9. It sometimes happens that a putative rearrangement coincides with an unsequenced gap in the reference genome, so it’s useful to visualize unsequenced gaps.

## Acknowledgements

We thank Takeshi Mizuguchi, Kazuharu Misawa, and Naomichi Matsumoto for helping us to fix inefficiencies in dnarrange.

## References

1. Hamada, M., Ono, Y., Asai, K., and Frith, M. C. (2017) Training alignment parameters for arbitrary sequencers with last-train Bioinformatics, 33(6), 926–928.

2. Frith, M. C. and Kawaguchi, R. (2015) Split-alignment of genomes finds orthologies more accurately Genome biology, 16(1), 1–17.

3. Huson, D. H., Albrecht, B., Ba?gci, C., Bessarab, I., Gorska, A., Jolic, D., and Williams, R. B. (2018) MEGAN-LR: new algorithms allow accurate binning and easy interactive exploration of metagenomic long reads and contigs Biology direct, 13(1), 1–17.

4. Frith, M. C. and Khan, S. (2018) A survey of localized sequence rearrangements in human DNA Nucleic acids research, 46(4), 1661–1673.

5. Mitsuhashi, S., Ohori, S., Katoh, K., Frith, M. C., and Matsumoto, N. (2020) A pipeline for complete characterization of complex germline rearrangements from long DNA reads Genome medicine, 12(1), 1–17.

6. Shafin, K., Pesout, T., Lorig-Roach, R., Haukness, M., Olsen, H. E., Bosworth, C., Armstrong, J., Tigyi, K., Maurer, N., Koren, S., et al. (2020) Nanopore sequencing and the Shasta toolkit enable efficient de novo assembly of eleven human genomes Nature biotechnology, 38(9), 1044–1053.

7. Shabardina, V., Kischka, T., Manske, F., Grundmann, N., Frith, M. C., Suzuki, Y., and Makalowski, W. (2019) NanoPipe—a web server for nanopore MinION sequencing data analysis GigaScience, 8(2), giy169.

8. Frith, M. C. (2011) A new repeat-masking method enables specific detection of homologous sequences Nucleic acids research, 39(4), e23–e23.

9. Grüning, B., Dale, R., Sjödin, A., Chapman, B. A., Rowe, J., Tomkins-Tinch, C. H., Valieris, R., and Köster, J. (2018) Bioconda: sustainable and comprehensive software distribution for the life sciences Nature methods, 15(7), 475–476.

10. Möller, S., Krabbenhöft, H. N., Tille, A., Paleino, D., Williams, A., Wolstencroft, K., Goble, C., Holland, R., Belhachemi, D., and Plessy, C. (2010) Community-driven computational biology with Debian Linux BMC bioinformatics, 11(Suppl 12), S5.

11. Morgulis, A., Gertz, E. M., Schäffer, A. A., and Agarwala, R. (2006) WindowMasker: window-based masker for sequenced genomes Bioinformatics, 22(2), 134–141.

12. Löytynoja, A. and Goldman, N. (2017) Short template switch events explain mutation clusters in the human genome Genome research, 27(6), 1039–1049.

13. Frith, M. C., Mitsuhashi, S., and Katoh, K. (2021) lamassemble: multiple alignment and consensus sequence of long reads In Multiple Sequence Alignment pp. 135–145 Springer

14. Lei, M., Liang, D., Yang, Y., Mitsuhashi, S., Katoh, K., Miyake, N., Frith, M. C., Wu, L., and Matsumoto, N. (2020) Long-read DNA sequencing fully characterized chromothripsis in a patient with Langer-Giedion syndrome and Cornelia de Lange syndrome-4 Journal of human genetics, 65(8), 667–674.

15. Frith, M. C., Nóe, L., and Kucherov, G. (2020) Minimally overlapping words for sequence similarity search Bioinformatics, 36(22-23), 5344–5350.

16. Mitsuhashi, S., Frith, M. C., Mizuguchi, T., Miyatake, S., Toyota, T., Adachi, H., Oma, Y., Kino, Y., Mitsuhashi, H., and Matsumoto, N. (2019) Tandem-genotypes: robust detection of tandem repeat expansions from long DNA reads Genome biology, 20(1), 1–17.

